# An Interscholastic Network to Generate LexA Enhancer Trap Lines in *Drosophila*

**DOI:** 10.1101/552513

**Authors:** Lutz Kockel, Catherine Griffin, Yaseen Ahmed, Lauren Fidelak, Arjun Rajan, Ethan P. Gould, Miles Haigney, Benjamin Ralston, Rex J. Tercek, Lara Galligani, Sagar Rao, Lutfi Huq, Hersh K. Bhargava, Ailis C. Dooner, Emily G. Lemmerman, Ruby F. Malusa, Tran H. Nguyen, Julie S. Chung, Sara M. Gregory, Kiyomasa M. Kuwana, Jonathan T. Regenold, Alexander Wei, Jake Ashton, Patrick Dickinson, Kate Martel, Connie Cai, Carissa Chen, Stephen Price, Jeffrey Qiao, David Shepley, Joanna Zhang, Meghana Chalasani, Khanh Nguyen, August Aalto, ByungJun Kim, Erik Tazawa-Goodchild, Amanda Sherwood, Ahmad Rahman, Sum Ying Celeste Wu, Joel Lotzkar, Serena Michaels, Hillary Aristotle, Antigone Clark, Grace Gasper, Evan Xiang, Frieda Luna Schlör, Melissa Lu, Kate Haering, Julia Friberg, Alyssa Kuwana, Jonathan Lee, Alan Liu, Emma Norton, Leena Hamad, Clara Lee, Dara Okeremi, Harry diTullio, Katherine Dumoulin, Sun Yu Gordon Chi, Grayson S. Derossi, Rose E. Horowitch, Elias C. Issa, Dan T. Le, Bryce C. Morales, Ayush Noori, Justin Shao, Sophia Cho, Mai N. Hoang, Ian M. Johnson, Katherine C. Lee, Maria Lee, Elizabeth A. Madamidola, Katrina E. Schmitt, Gabriel Byan, Taeyoung Park, Jonathan Chen, Alexi Monovoukas, Madison J. Kang, Tanner McGowan, Joseph J. Walewski, Brennan Simon, Sophia J. Zu, Gregory P. Miller, Kate B. Fitzpatrick, Nicole Lantz, Elizabeth Fox, Jeanette Collette, Richard Kurtz, Chris Duncan, Ryan Palmer, Cheryl Rotondo, Eric Janicki, Townley Chisholm, Anne Rankin, Sangbin Park, Seung K. Kim

## Abstract

Binary expression systems like the LexA-LexAop system provide a powerful experimental tool kit to study gene and tissue function in developmental biology, neurobiology and physiology. However, the number of well-defined LexA enhancer trap insertions remains limited. In this study, we present the molecular characterization and initial tissue expression analysis of nearly 100 novel StanEx LexA enhancer traps, derived from the *StonEx^1^* index line. This includes 76 insertions into novel, distinct gene loci not previously associated with enhancer traps or targeted LexA constructs. Additionally, our studies revealed evidence for selective transposase-dependent replacement of a previously-undetected *KP* element on chromosome III within the StanEx^1^ genetic background during hybrid dysgenesis, suggesting a molecular basis for the over-representation of LexA insertions at the *NK7.1* locus in our screen. Production and characterization of novel fly lines were performed by students and teachers in experiment-based genetics classes within a geographically diverse network of public and independent high schools. Thus, unique partnerships between secondary schools and university-based programs have produced and characterized novel genetic and molecular resources in *Drosophila* for open-source distribution, and provide paradigms for development of science education through experience-based pedagogy.

## Introduction

Binary gene expression systems are an important foundation for investigating and manipulating *Drosophila* gene expression with temporal and cellular specificity. Generation of a yeast GAL4-based transactivator to induce expression of target genes fused to GAL4-responsive upstream activating sequences (UAS), has established a widely-used binary gene expression system in *Drosophila* (Brand & Perrimon 1993; Hayashi et al., 2002; Gohl et al., 2011). Common examples of GAL4-based gene fusions include transgenes with relatively short enhancer elements that direct GAL4 expression, and endogenous enhancer-directed GAL4 expression following random genome insertion by transposons encoding *GAL4* (‘enhancer trapping’ O’Kane & Gehring, 1987).

Studies of many biological problems benefit from simultaneous manipulation of two or more independent cell populations or genes (Rajan & Perrimon 2011). In prior studies, parallel use of two binary expression systems allowed insightful clonal analysis of multiple cell populations (Lai & Lee 2006; Bosch 2015), powerful studies of epistasis between different tissues (Yagi et al., 2010; Shim et al., 2013), and discovery of specific cell-cell contacts (Gordon & Scott 2009; Bosch et al., 2015; Macpherson et al. 2015). This multiplex approach requires an additional expression system that functions independently of UAS-Gal4 system, like the LexA system derived from a bacterial DNA binding domain (Szűts & Bienz 2000; Lai & Lee 2006; Pfeiffer et al., 2010; Knapp et al., 2015; Gnerer et al. 2015). In that system, fusion of the LexA DNA binding domain to a transactivator domain generates a protein that regulates expression of transgenes linked to a LexA operator-promoter (LexAop). Unfortunately, the number and quality of fly lines expressing a LexA transactivator remains small compared to the thousands of comparable GAL4-based lines. From a scholastic network including Stanford University and secondary school science classes in New Hampshire, New York, New Jersey and Illinois we generated novel LexA-based enhancer trap drivers for fly biology.

## Material and Methods

Fly husbandry, intercrosses, immuno-histochemistry and molecular biology were performed as previously described in (Kockel et al 2016).

### Immuno-histochemistry (IHC)

L3 larvae from cross of *StanEx^novel msertlon^* with a line harboring a LexA operator- GFP reporter transgene (*LexAop2-CD8::GFP*; Pfeiffer et al., 2010) were dissected in PBS and fixed in 4% Formaldehyde/PBS for 30 mins, permeabelized in 0.2% Triton X-100/PBS for 4 hrs, and blocked in 3% BSA Fraction V/PBS for 1 hr. Incubation of primary and secondary antibodies were O/N in 3%BSA/PBS at 4° Celsius using a platform rocker. All specimens were rinsed (1 min) and washed (20 mins) three times with PBS after antibody incubations. Primary Antibody: Goat anti-GFP 1:3000 (Rockland 600-101-215). Secondary antibody: Donkey antiGoat Alexa488 (Life Technologies, A11055). All samples were mounted in SlowFade Gold mounting medium with DAPI (Life Technologies, S36938).

### Microscopy

Microscopy was performed on a Zeiss Axiolmager M2 with Zeiss filter sets 49 (DAPI) and 38HE (Alexa488) using the extended focus function.

### Fly husbandry and fly strains

A standard cornmeal-molasses diet was used to maintain all fly strains (http://flystocks.bio.indiana.edu/FlyWork/media-recipes/molassesfood.htm). The following strains were used: w[*]; ry[506] Sb[l] Ρ{ry[+t7.2]=Δ2-3}99Β/ΤΜ2, Ubx (Bloomington 1798), w[*]; P{y[+t7.7] w[+mC]=26XLexAop2-mCD8::GFP}attP2 (Bloomington 32207), y[l],w[1118]; P{w[mC]=LHG]StanEx[l]} (Bloomington 66673), w[*]; L[*]/CyO; ftz[*] e[*]/TM6,Tb[*].

### Hybrid dysgenesis

F_0_: Females of donor stock y,w,StanEx[l] were mated to males w[*]; ry[506],Sb[l]Δ2-3/TM2,Ubx. F_1_: y,w,StanEx[1]; ry[506],Sb[l], Δ2-3/+ males were crossed to w[*]; L[*]/CyO; ftz[*] e[*]/TM6,Tb,Hu females. F_2_: *w+* males were mated to w[*]; L[*]/CyO; ftz[*] e[*]/TM6,Tb,Hu. F_3_: The insertion line was stably balanced deploying a brother-sister cross.

This P-element vector also enables subsequent recombinase-mediated cassette exchange (RMCE; Gohl et al. 2011).

### Insertion site cloning

We followed an inverse PCR approach (Kockel et al, 2016, http://www.fruitfly.org/about/methods/inverse.pcr.html), to molecularly clone the insertion sites of StanEx P-elements. DNA restriction enzymes used: Sau3AI (NEB R0169) and Hpall, (NEB R0171). Ligase used: T4 DNA Ligase (NEB M0202). 5’ end cloning: Inverse PCR primer “Plac1” CAC CCA AGG CTC TGC TCC CAC AAT and “Plac4” ACT GTG CGT TAG GTC CTG TTC ATT GTT. 3’ end cloning: Primer pair “Kurt” TGT CCG TGG GGT TTG AAT TAA C and “Ulf” AAT ACT ATT CCT TTC ACT CGC ACT. Sequencing primer 5’ end: “Sp1” ACA CAA CCT TTC CTC TCA ACA. Sequencing primer 3’ end: “Ulf” or “Berta” AAG TGG ATG TCT CTT GCC GA. For insertions where the sequence of one end only could be determined by inverse PCR, we pursued a gene-specific PCR approach (Ballinger and Benzer 1989) using P-element and gene-specific primers. 5*′* end specific P-element primer “Chris”: GCA CAC AAC CTT TCC TCT CAA C, sequencing primer 5’ end: “SP1”. 3’ end specific P-element primer “Dove”: CCA CGG ACA TGC TAA GGG TTA A, sequencing primer 3’ end: “Dove”. Sequence of gene-specific primers are available upon request.

### Generation of Sequence Logos and position frequency matrices (PFMs)

The generation of sequence logos was performed as described (Crooks et al., 2004) using the web tool http://weblogo.threeplusone.com/. The input sequence motif data is listed in **Suppl. Table 2.** The insertion site sequence is displayed and utilized with the genomic scaffold co-directionally oriented to inserted P-element (Linheiro and Bergman, 2008). If P-elements are inserted 5’->3’, the strand of insertion was called +, and unprocessed genomic scaffold sequences were used to extract the insertion site sequences. If P-elements are inserted 3’->5’, the strand of insertion is called -, and the reverse complement of the genomic scaffold sequences were used to extract these insertion site sequences.

Position frequency matrices were generated using data from (Linheiro and Bergman, 2008), (Gohl et al, 2011) and (Kockel et al., 2016). For GT P-element PFMs, quality issues of the insertion site data was noted (Linheiro and Bergman, 2008), and only the insertions on the + strand were used (Linheiro and Bergman, 2008). As a result of only 12 InSite P-elements insertion sites sufficiently mapped (Gohl et al., 2011), the PFM was not testable (0% C and 0% G in position 1; 0% G at position 13). All data is displayed in **Suppl. Table 3.** Chi-square testing was performed in MS Excel (2007) using the CHITEST function.

### Sequencing the genomic site in NK7.1 / Heatr2 around hot spot at 3R:14,356,561

The genomic site was amplified in 5 fragments of overlapping segments of approx. 500bp called A, B, C, D, and E Fragments. C Fragment contained the hot spot 3R:14,356,561. Co-ordinates A-Fragment: 3R:14,355,462 - 3R:14,355,962. Primer sequences for A-Fragment amplification: NK7.l-GS_A.FOR: AGTGGAAACGAGCGAAGCTG; NK7.1-GS_A.REV: AAAGGTCAAATGTGATGCAGCGAG. Co-ordinates B-Fragment 3R:14,355,862 - 3R:14,356,362. Primer sequences for B-Fragment amplification: NK7.1-GS_B.F0R: TCTGTGCAGATAGGAAATTACTCATT and NK7.1-GS_B.REV: CTCTTGCCACTTTCTGTGAGCTT. Co-ordinates C-Fragment: 3R:14,356,190 - 3R:14,356,709. Primer sequences for C-Fragment amplification: NK7.1-GS_C.FOR2: CTGGGCCAGTCAAGTGTGTA and NK7.1-GS_C.REV2: AGAGCTACGAACCTGGCC. Co-ordinates D-Fragment: 3R:14,356,662 - 3R:14,357,162. Primer sequences for D-Fragment amplification: NK7.1-GS_D.F0R: GCGATGAGGATGAAGTTGTCGG and NK7.1-GS_D.REV: GACTCTCTTCATCGCCAGCC. Co-ordinates E-Fragment: 3R:14,357,124 - 3R:14,357,638. Primer sequences for E-Fragment amplification: NK7.1-GS_E.F0R: CCTGGCCATAGAGATCCAAG and NK7.1-GS_E.REV: TGCGAAGCTGCAAAGTAAAA. According to the published genomic sequence, the expected size of the C-Fragment is 520 bp. PCR in genomic StanEx1 DNA amplified a DNA fragment of 1682bp, consisting of C-Fragment sequence containing KP element sequence.

All primer and genomic reference sequences are also deposited in **Suppl. Table 4,** Workbook “Genomic Seq Flybase”.

### Sequencing KP element at 88B4-6

The KP element at 88B4-6 of the StanEx^1^ strain was amplified using the C-Fragment primer (see above) NK7.1-GS_C.FOR2: CTGGGCCAGTCAAGTGTGTA and NK7.1-GS C.REV2: AGAGCTACGAACCTGGCC. Due to the A:T-rich region and the resulting A:T stutter during sequencing 5’ of the insertion site, all sequencing was performed on the lagging strand, 3’->5’ relative to the genomic scaffold in three replicates. Sequencing primers: NK7.1-GS_C.REV2 (see above), NK7.1-KP_BPS_REV1: TAGGTACGGCATCTGCGTTG, KP_BPS_REV2: CAGCCTTCCACTGCGAATCAKP_BPS_REV3: CAAGGCTCTGCTCCCACAAT. All individual sequence and primer data is shown in **Suppl. Table 4,** Workbooks: “Seq StanEx KP Element” and “Primer Sequences”.

### Testing Strains for KP element insertions

Genomic DNA was extracted using standard methods and presence of KP element(s) was tested using Primers NK7.1-KP_3_REV2, ATGCCCAGGATGAATTGAAA and NK7.1-KP_3_FOR2, TCAGATGTGGAAACGTCGAT. The following strains were tested for the presence of KP element(s) in their genome: *Oregon R* (Bloomington Stock #5); *bw^1^* (Bloomington Stock #245); *cn^1^* (Bloomington Stock #263); *y^1^,w^1^* (Bloomington Stock #1495); *w^1^* (Bloomington Stock #2390); *w^1118^* (Bloomington Stock #5905); *w^1118^* iso II, iso III (Bloomington Stock #6326); Oregon-R-SNPiso3 (Bloomington Stock #6363); Canton-S-SNPiso3 (Bloomington Stock #6366); Canton-S (Bloomington Stock #64349). Only *w^1^* (Bloomington Stock #2390) was found KP positive.

### GenBank accession for KP element

The GenBank accession number for the KP element sequence at 88B4-6 characterized in this study is MK510925. The 5’->3’ annotated KP element sequence is also displayed in **Suppl. Table 4,** Workbook “Reconstituted StanEx-KP seq 5-3”.

### Probability calculation of StanEx P-element insertion site hot spot at 3R:14,356,561, 88B4-6

We calculated the probability that a single genomic P-element insertion site would be selected at least 9 times in a series of 188 (total number of StanEx insertions generated so far) random insertions. The number of confirmed and non-identical individual P-element insertion sites present on the autosomes of the genome of *Drosophila melanogaster* was determined conservatively by counting confirmed and unique EPgy2, GT1, SUPor-P, GawB(+) and XP insertions (Linheiro and Bergman, 2008). Hence, we tested the null hypothesis that the transposable element is equally likely to insert itself at any of 8,161 target sites. Mathematically, *p* = Pr (*X*_1_ ≥ 9 ∪ *X_2_* ≥ 9U*X*_3_ ≥ 9 ∪ … ∪ *X*_8161_ ≥ 9), where Pr is given by a multinomial distribution with all event probabilities equal to 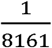 and 188 trials. Since the computation of the exact p-Value is computationally expensive, we approximated the p-Value by using the simplification that Pr(*X*_1_ ≥ 9 ∪ *X*_2_ ≥ 9 ∪ *X*_3_ ≥ 9 ∪ … ∪*X*_8161_ ≥ 9) ≤ Pr(*X*_1_ ≥ 9) + Pr(*X*_2_ ≥ 9) + … + Pr(*X*_8161_ ≥ 9) = 8161·Pr(*X*_1_ ≥ 9). The value of Pr (*X*_1_ ≥ 9) can be calculated using a binomial distribution with success probability and 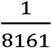 188 trials. Thus, we calculated the value of 8161 • Pr (*X*_1_ ≥ 9) using the R command: 8161 * pbinom(q = 8, size=188, p = 1/8161, lower.tail=FALSE).

The probability of obtaining nine insertions into an identical genomic site by chance is estimated to be small (*P* < 3.32 x 10^-17^). For the purpose of comparison, the chances to win the California Power Ball Lottery in October 2018 were 2.9 x 10^-8^. To corroborate this result empirically, we ran a computer simulation of the stochastic process in which a transposable element was inserted into an array of 8,161 equally probable target sites 188 times. After 100,000 runs of the simulation, we did not detect choice of the same insertion site 9 times.

### Secondary school class descriptions

We formed partnerships between Stanford University investigators and classes at 4 high schools in the U.S. The class at Phillips Exeter Academy (NH) was taught as two 11-week courses “BIO590” and “BIO670” in the winter and spring terms. One year classes called "Research 11/12" ran at Commack High School (NY), “BIO670” at Pritzker College Prep (Chicago, IL), and “Sci574” 2/3 class + 1/3 elective extension at The Lawrenceville School (NJ). Pre-requisites for admission to these class were determined by individual schools, and included no pre-requisites (Commack High School and Pritzker College Prep), advanced placement (AP) biology or one term of a genetics elective (Phillips Exeter Academy, NH), or one term advanced science (The Lawrenceville School).

Bio670 / Res 11/12 / Sci574 were scheduled for three to four 50 minute periods / week, and up to 5-6 unscheduled hours per week. The instruction manuals, additional manuals for teachers, schedules and related problem sets are available on request.

At Exeter, students spent about nine weeks executing the hybrid dysgenesis crosses including mapping and balancing of novel strains. Further characterization of the insertion site was performed by polymerase-chain reaction and DNA sequencing using standard genomic DNA recovery. Crosses with reporter strains (*LexAop2-CD8::GFP*) were performed during the final 3 weeks of class, permitting instruction in larval dissection and microscopy to document tissue expression patterns of candidate enhancer traps. This schedule was modified to fit the year-long schedule at other secondary schools. Based on class performance and teachers’ evaluations, selected students were invited to continue studies in the Kim group at Stanford University School of Medicine during summer internships lasting about 6 weeks. These studies included further molecular mapping of transposon insertion sites, and verification of tissue patterns of enhancer trap expression. Students returning in the fall term helped instructors to run the subsequent iteration of Bio670, and also pursued independent projects.

### Data and reagent availability

All StanEx derivatives and associated data are available at the Bloomington stock center. All molecular and image data are additionally available at http://stanex.stanford.edu/about/.

## Results

### Generating novel LexA-based enhancer trap strains

To generate LexA-based enhancer trap fly lines, we mobilized the P-element in a previously characterized *StanEx^1^* line (Kockel et al 2016). The *StanEx^1^* strain contains a single X-linked derivative of the InSITE P-element (Gohl et al., 2011) harboring a weak P-promoter driven cDNA encoding a LexA DNA-binding domain fused to the hinge-transactivation domain of Gal4 (LexA::HG, Yagi et al., 2010). We mobilized this X-linked *StanEx*^1^ *P*-element to the autosomes using transposase Δ2-3 at 99B (Robertson et al., 1988), to generate LexA P-element enhancer trap lines using standard methods **(Methods; Suppl. Table 1;** O’Kane et al., 1987). Our goal was to permit interaction of the weak promoter in the mobilized *StanEx*^1^ *P*-element with the local enhancer environment of the insertion site, and thereby generate spatial and temporal expression specificity of each LexA::HG insertion (O’Kane and Gehring, 1987).

### Mapping StanEx *P*-element insertion sites

We next used established inverse PCR-based molecular methods to map the chromosomal insertion position of the *StanEx*^1^ *P*-elements to the molecular coordinates of the genomic scaffold **(Figure 1;** http://stanex.stanford.edu/about/). The 93 novel insertions of this study were equally distributed across autosomes II and III, and their chromosomal arms (2L, 24 insertions: 2R, 24 insertions: 3L, 20 insertions: 3R, 25 insertions). In this collection, we included 2 lines (SE133 and SE174) that inserted into repetitive DNA, and whose insertion site we could not map molecularly. We excluded 7 lines that were inserted into the identical location in *NK7.1/Heatr2* at 3R:14,356,561, as this precise location was tagged by prior StanEx insertions (see below, lines RJ-3 and EH-4; Kockel et al., 2016). As observed previously (Bellen et al, 2011), the majority of novel insertions (81/91 or 89%) integrated into 5’ gene elements, including promoters, and first exons or introns. Of the 93 novel insertions presented here, we observed an even distribution of insertional direction by the *StanEx*^1^ *P*-element into genomic DNA. Using the 5’ and 3’ end of the *P*-element as coordinates, we found 46/93 insertions were oriented 5’->3’, and 44/93 insertions were oriented 3’->5’. In three cases we were unable to determine the direction of P-element insertion. In two of these three cases, the *StanEx*^1^ *P*-element inserted into repetitive DNA (see above). In one of these three cases (SE444) the *StanEx*^1^ *P*-element inserted into the Hsp70 locus at 87A2 that consists of a 1:1 mirror-image arrangement of two nearly-identical copies of the hsp70 promoter and coding region (Hsp70Aa, FBgn0013275 and Hsp70Ab, FBgn0013276); sequence analysis of 5’ and 3’ adjacent genomic DNA in this line failed to unambiguously resolve the orientation of the SE444 *P*-element **(Suppl. Table 1).**

**Figure 1:**
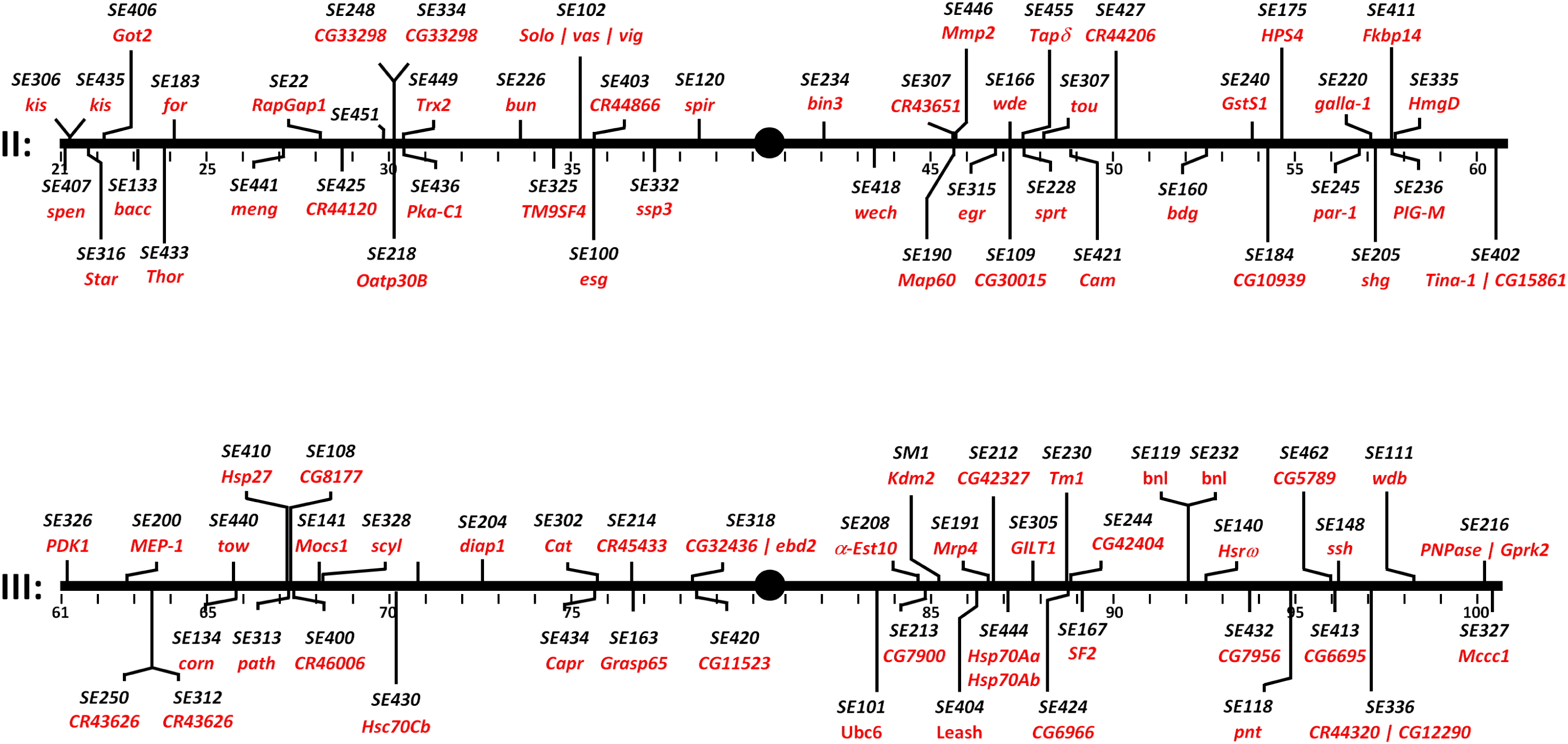
Insertion sites and genes tagged by LexA::HG enhancer traps on autosomes II and III. Associated molecular data is detailed in **Suppl. Table 1.**

We then analyzed the number of novel loci tagged by our *StanEx*^1^ LexA enhancer trap. To search for genes tagged by multiple constructs or insertions, we surveyed publically-available lines cataloged in FlyBase, including previous StanEx releases, and lines generated by the FlyLight project (Pfeiffer et al, 2013). 76 of 91 mapped lines presented here mark novel loci, among them *eiger* (SE315), *Star* (SE316), *Thor* (SE433), *kismet* (SE306 and SE435), *par-1* (SE245), and *branchless* (SE 119 and SE232). Of note, we obtained insertions into several long non-coding RNA loci (lncRNA:CR43626, SE250 and SE312; lncRNA:CR43651, SE307; lncRNA:CR44120, SE425; lncRNA:CR44206, SE427; lncRNA:CR44320, SE336; lncRNA:CR45433, SE214; and lncRNA:CR46006, SE400). Apart from transcriptional mapping (Graveley et al., 2010), these non-coding RNAs have not been functionally characterized.

Of the fifteen genes previously tagged by LexA, four were tagged in FlyLight lines (*meng-po,* SE441, *bunched,* SE226, *G-coupled Receptor Kinase 2,* SE216, *pointed,* SE118), ten were identified by prior StanEx insertions (*RapGapl* SE422, *split ends* SE407, *incRNA:CR43626* SE312 and SE250, *bicoid interacting protein 3* SE234, *α-Esterase-10* SE208, *Hsrω* SE140, *bacchus* SE133, *escargot* SE100, and the *solo, vasa, vig* locus SE102), and one was tagged in both FlyLight and StanEx studies (SE120, *spire*; see **Suppl. Table 1).** In summary, our approach generated multiple novel LexA-based autosomal enhancer traps.

### Selected tissue expression of LexA in the StanEx collection

To verify the use of the *StanEx*^1^ P-element for enhancer trapping, we intercrossed this line with flies harboring a transgene encoding a LexA operator linked to a cDNA encoding a membrane-GFP reporter *(iexAop2-CD8::GFP*; Pfeiffer et al., 2010) and confirmed membrane-associated GFP expression in several tissues, including larval and adult tissues (data not shown). Next we used this strategy to assess the tissue expression patterns of novel insertion lines. 3^rd^ instar larvae of bi-transgenic offspring were analyzed by immuno-histochemical (IHC) staining for GFP expression, and simultaneous counter-staining for cell nuclei (DAPI). Image data from selected LexA enhancer trap lines were collected and tissue expression catalogued **(Suppl. Table 1).** Within the collection, we detected GFP expression in nearly all tissues of the L3 larva, including a variety of neuronal cell types in the Central Nervous System (CNS), Ventral Nerve Cord (VNC: **Figure 2)** and Peripheral Nervous System (PNS), imaginai discs, and a wide range of other somatic tissues like fat body, malphigian tubules and trachea **(Suppl. Table 1).** In the cases of lexAop-CD8::GFP expression directed by LexA from an insertion in the *solo/vasa/vig* locus (SE102), *Diap1, a-Esterase 10* (SE208), *OatP30B* (SE218), *NK7.1* (SE229) and *cornetto* (SE134), we observed distinct patterns of cell labeling in the CNS, VNC and ring gland **(Figure 2A-F).** In *Diap1-LexA; lexAop-CD8::GFP* larva we noted strong staining of neurons in the *pars intercerebralis* of the CNS, corresponding to neuroendocrine insulin-producing cells (IPCs: **Figure 2B).**

**Figure 2:**
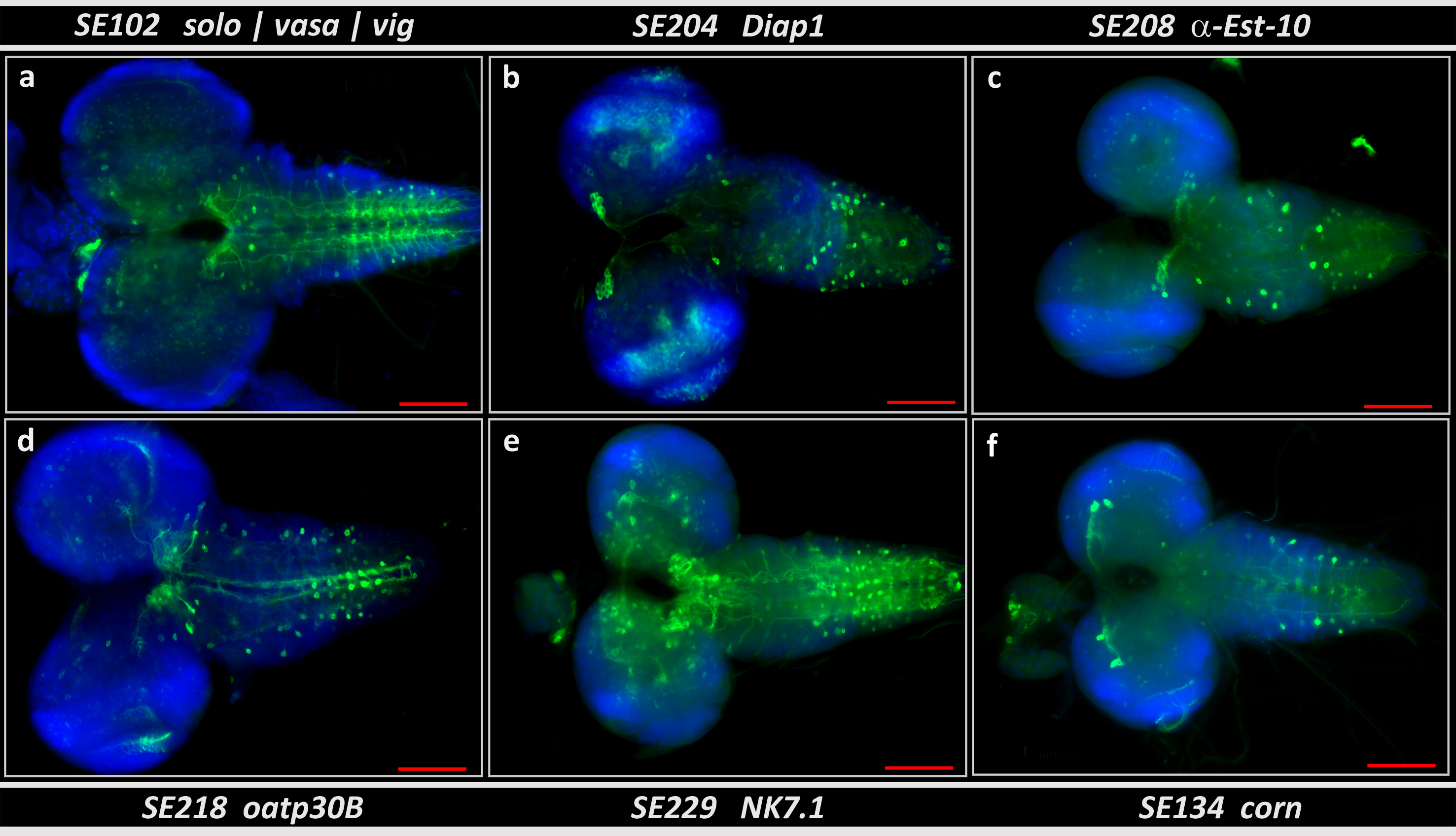
Expression pattern of six selected StanEx enhancer trap insertions in central nervous system (CNS), ventral nerve chord (VNC) and ring gland (RG) complexes of third instar larvae. Enhancer traps of (a) SE102 (insertion in *solo/vasa/vig),* (b) SE204 (insertion in *Diap1),* (c) SE208 (insertion in *α-Est-10),* (d) SE218 (insertion in *oatp30B),* (e) SE229 (insertion in *NK7.1)* and (f) SE134 (insertion in *corn)* crossed to *lexAop-CD8::GFP* are shown. Green: anti-GFP, Blue: DAPI. Scale bar = 100 μm.

To facilitate accessibility of the molecular and image data **(Suppl. Table 1),** we uploaded these to the searchable StanEx website (http://stanex.stanford.edu/about/; Kockel et al 2016), a database searchable by expression pattern, cytology and specific genes. This includes supplementary image analysis, data from immunostaining and molecular features of *StanEx*^1^ insertion loci, and is freely accessible to the scientific community.

### An unrecognized KP Element in the StanEx^1^ line

During the generation of 188 individual StanEx enhancer trap P-element lines (from this work and Kockel et al 2016), we observed nine independent insertions into the *Nk7.1* / *Heatr2* locus at 88B4-6 **(Figure 3).** Molecular characterization of these nine insertions revealed 3R:14,356,561 as the common insertion coordinate. This particular insertion hot spot was not reported in prior hybrid dysgenesis efforts, which included use of a variety of distinct *P*-element constructs (Bellen et al., 2011, Linheiro and Bergman, 2008). The probability of obtaining 9/188 insertions into an identical genomic site by chance is small (estimated *P* < 3.32 × 10^-17^: see Methods).

**Figure 3:**
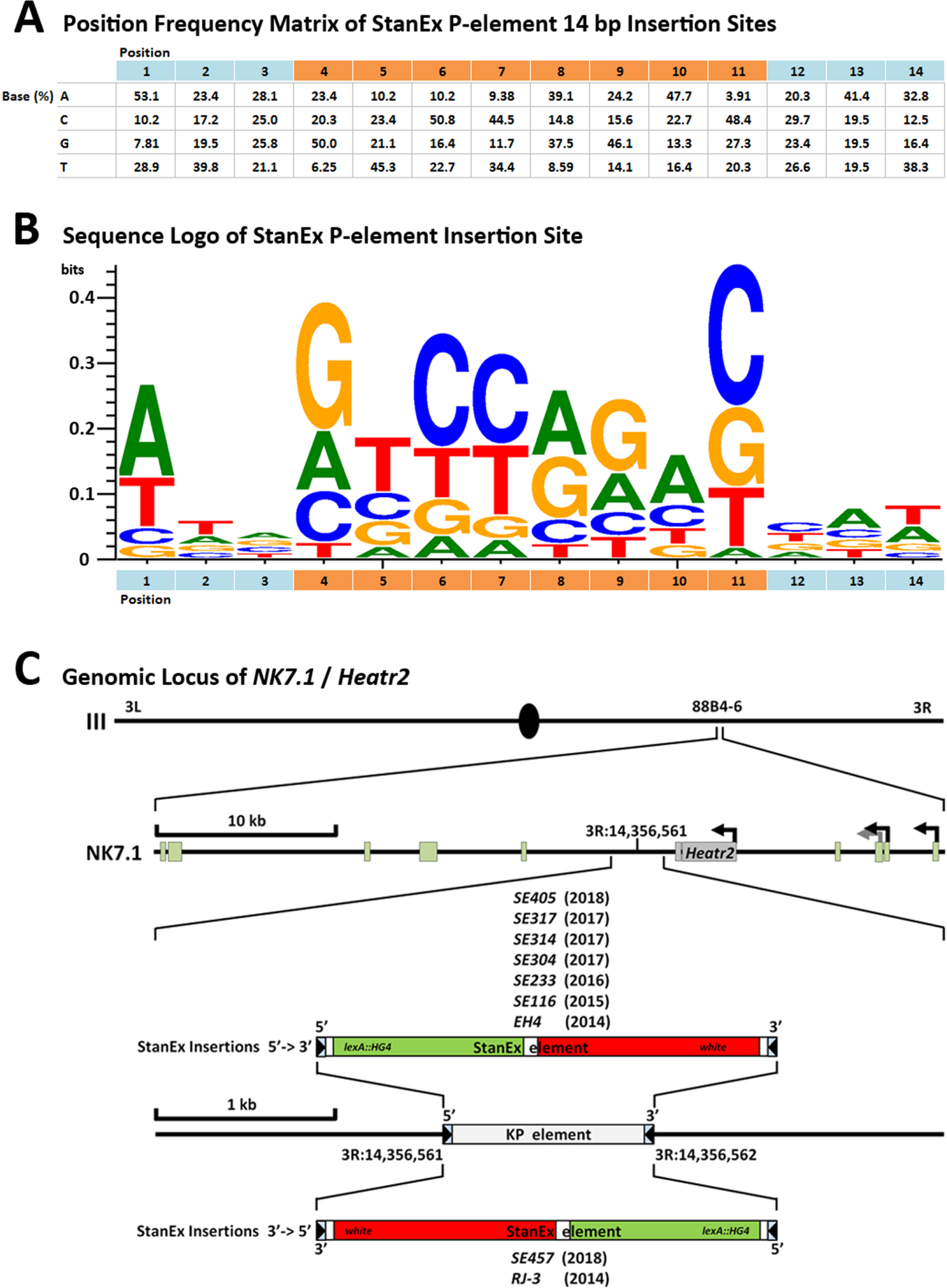
Properties of the StanEx enhancer trap *P*-element. (A) Position frequency matrix (PFM) of the 14 bp *P*-element insertion site (see text for details, **Suppl. Table 2).** Base position 1-14 in 5’->3’ direction on X-axis. The 8 bp sequence that will give rise to the directed repeat *after* insertion is highlighted in orange. Base composition (in %) on Y-axis as indicated. (B) Sequence logo derived from (A), X-axis as in (A), Y-axis in bits (Crooks et al., 2004). (C) Insertion site of 9 StanEx lines within the locus of *NK7.1/Heatr2* at 3R:14,356,562 88B4-6 on chromosome III. Transcription site start arrows mark alternative first exons of NK7.1. SE-number identifiers of StanEx enhancer traps and the year of their derivation are shown along their insertion direction **(Suppl. Table 4)**.

The presence of this observed *P*-element insertion hotspot suggested a non-classical mechanism of targeting or insertion of the *StanEx*^1^ element. Preferential target site selection of *P*-elements based on the presence of DNA homology between the P-element and its target site has been described (Taillebourg and Dura, 1999). To investigate a possible homology-based mechanism targeting the insertion into the hotspot at 88B4-6, we searched for apparent DNA homology between the *StanEx*^1^ enhancer trap *P*-element and the published *D. melanogaster* genomic sequences present in Release 6 of Flybase (Hoskins et al., 2015) at that site, and did not detect significant homologies. Hence, we explored the possibility of an insertion site bias inherent to our *StanEx*^1^ P-element construct. We generated a position frequency matrix (PFM, **Figure 3A)** and sequence logo of the 14 bp P-element insertion site motif (Majumdar and Rio, 2014) using 128 unique *StanEx*^1^ insertions that were characterized by insertion site sequencing with single nucleotide precision on the 5*′ and* 3’ ends (Crooks et al., 2004; **Figure 3B).** This *StanEx*^1^ PFM was individually compared to PFMs of the P elements EPgy2 (n=3112), GT1 (n=465), SUPor-P (n=2009), GawB(+) (n=1072) and XP (n=4126) **(Suppl. Table 3;** Linheiro and Bergman, 2008). We also compared the *StanEx*^1^ PFM to a ‘summary’ consensus motif based on available P-element insertion site data **(Suppl. Table 3).** Using χ^2^ testing, we were unable to detect a significant difference of the *StanEx*^1^ insertion site motif and any of the other individual or agglomerated P-element insertion site motifs **(Suppl. Table 3).** We conclude that *StanEx*^1^ and other P-elements have similar insertion site preferences.

Thus, we assessed the possibility that the specific 3R:14,356,561 target site harbored by the *StanEx*^1^ starter strain might contain sequences not annotated in the published *D. melanogaster* genome (Version 6), and that might contribute to the modestly increased targeting frequency at this locus. Sequencing the *StanEx*^1^ *P*-element recipient hotspot at 3R:14,356,561 in our *StanEx*^1^ donor strain, prior to hybrid dysgenesis, revealed the presence of an unrecognized 1.1 kb *KP* element at 3R:14,356,561 **(Figure 3C, Suppl. Table 4,** GenBank Accession MK510925). This *KP* element was flanked by the 8 bp target site direct duplication GCCCAACC. A *KP* element is a non-autonomous *P*-element with intact inverted repeats, whose transposase-encoding exons 2-4 contain deletions. These deletions produce an (ORF) frame that encodes a type II repressor instead of functional transposase (Black et al., 1987; Rio, 1990; Majumdar and Rio, 2014; Kellerher, 2016), permitting *KP* element mobilization only when transposase is provided in *trans.* Thus, an interaction of a KP-element with the StanEx *P*-element in the *StanEx*^1^ starter strain could underlie the observed repeated integration of StanEx P-element into 88B4-6.

To investigate whether the *KP* element was deleted upon *StanEx*^1^ insertion at 3R:14,356,561, we analyzed DNA sequence generated by inverse and conventional PCR covering the breakpoint between the StanEx P-element and adjacent genomic sequences in these **9** StanEx lines. We also attempted to amplify *KP* specific sequences using *KP*-specific primers. In the nine insertions of *StanEx*^1^ into 3R:14,356,561, no *KP* element sequences were detected (data not shown). Thus, in the process of the hybrid dysgeneses that gave rise to *StanEx*^1^ insertions at 3R:14,356,561, the *KP* element was concurrently deleted at that site. Additional molecular analysis **(Fig 3A-C, Suppl. Table 4)** revealed that in **7/9** cases, this KP replacement by the *StanEx*^1^ P-element conserved the direction of the original KP-element, and in **2/9** cases the P-element replacement led to small genomic DNA deletions adjacent and 5’ to the integration site **(Suppl. Results 1** and **Suppl. Table 4).** Together these findings suggest that the *StanEx P*-element replaces the *KP* element at the site 3R:14,356,561 in all cases.

### Creating an interscholastic network to generate resources for Drosophila genetics

In a prior study, we produced and characterized novel fly enhancer trap lines through a unique course partnering students and instructors in a genetics class (Bio670) at an independent New Hampshire secondary school (Phillips Exeter Academy) with university-based researchers (Kockel et al., 2016). To test if this paradigm could be expanded to include additional classes, we developed a second molecular biology class at Exeter that mapped *StanEx*^1^ P-element genomic insertions with inverse PCR-base molecular methods (Bio590). The Bio590 class was taught by teachers who also led Bio670. This expansion of the curriculum to include experimental molecular biology relieved a bottleneck that arose due to the single term duration (11 weeks) of Bio670, that left >50% of newly-generated *StanEx*^1^ insertion lines unmapped. Thus the two classes, Bio590 and Bio670, integrated and enhanced the longitudinal quality of genetic experiments presented here.

We next assessed if the curriculum of fly-based transmission genetics and hybrid dysgenesis, molecular characterization of insertion lines, and expression analysis with the *iexAop-GFP* reporter gene could be adapted to year-long genetics classes at other secondary schools. Over a 4 year span (2016-2019), we established our curriculum at high schools in New York (Resll/12, Commack High School), Illinois (Bio670, Pritzker College Prep), and New Jersey (Sc574, The Lawrenceville School; **Figure 4).** Thus, data and resources detailed here stem from secondary schools collaborating throughout the academic year with a research university (Stanford). To foster production and sharing of data and fly strains, and to achieve student learning goals, the partners in this interscholastic network benefitted from structured interactions, including (1) summer internships for students (n=17) or instructors (n=9) with the university research partner, (2) weekly term-time research teleconferences organized by university partners with high school instructors and classes, and (3) annual site visits of university collaborators to secondary school classes during term time **(Figure 4).** Multiple initially un-programmed pedagogical outcomes resulted for students and teachers at partnering schools. These included service by students, who completed these classes, as proctors or teaching assistants in the next term (n=10), instruction of incoming teachers by students who successfully completed the course in a prior year (n=3), and collaboration between science instructors at different schools to establish new science curriculum through direct consultation and sharing of open-source materials (n=4). Thus, a consortium of students, teachers and leadership at multiple, geographically-unconnected secondary schools and university-based programs have formed a unique research network actively generating well-characterized fly strains suitable for investigations by the community of science.

**Figure 4:**
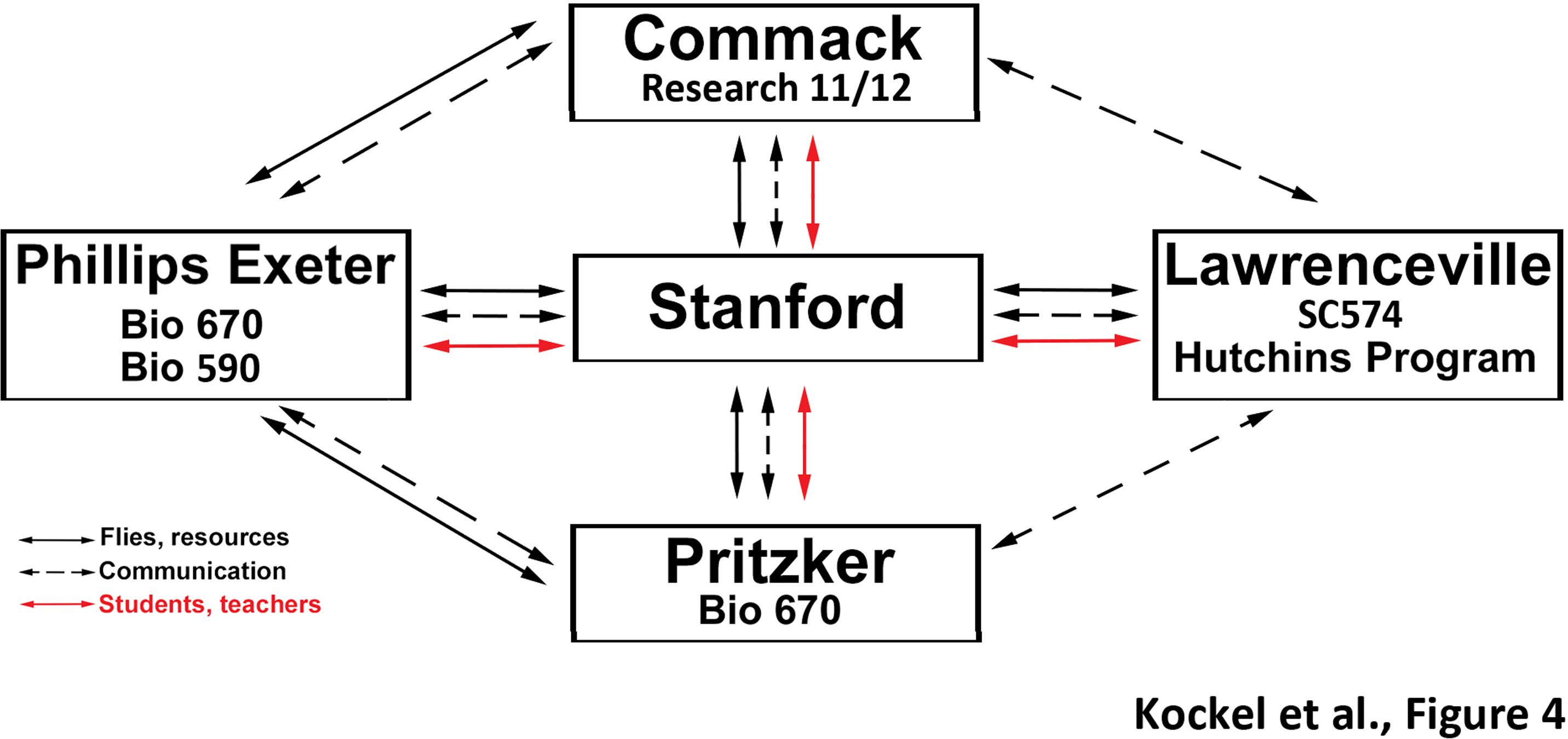
Structure of scholastic network and routes of resource exchange. The name of the high school and class number specification is shown. The exchange of materials and other resources (black contiguous arrows), student, teacher and instructor visits (red arrows) and voice/video/email communication (black dotted arrows) is shown.

## Discussion

Here we used P-element mobilization in *Drosophila melanogaster* to generate 96 new fly lines tagging 76 novel loci with a LexA enhancer trap. Expression of LexAop-based reporters suggests that enhancer traps in this collection are distinct, and are expressed and active in a wide variety of tissues, including neurons within the CNS and VNC. This collection has been submitted to an international fruit fly repository (Bloomington Stock Center) and should prove useful for genetic, developmental and other studies of cells and tissues. Assessment of an apparently biased insertion frequency at locus 88B4-6 in our screen led to discovery of a previously undetected *KP* element in the *StanEx*^1^ starter strain, and we present evidence for the *StanEx*^1^ P-element replacing the *KP* element at this locus, a finding that will influence future enhancer trap screens. Data and biological resources here were generated from partnerships connecting a research university with teachers and students at multiple secondary schools. This illustrates the feasibility of building an interscholastic network to conduct biological research that impacts the community of science, and provides an ‘experiential’ paradigm for STEM education.

Experimental approaches in biology benefit from temporal-or cell-specific control of gene expression, like that possible with binary expression strategies pioneered in the *Drosophila* GAL4-UAS system (Brand and Perrimon, 1993). Intersectional approaches, like simultaneous use of the LexA-lexAop and GAL4-UAS systems, have also greatly enhanced experimental and interpretive power in fly biology, particularly studies of neuroscience and intercellular communication (Simpson, 2016, Martin et al., 2017, Dolan et al., 2017). For example, enhancer traps generated here **(Suppl. Table 1)** include insertions into genes regulating: (1) the Insulin Receptor-Akt-TORC1 pathway, like *Pdk1, Scylla* (ortholog of the HIF-1 target REDD1), *widerborst* (ortholog of PP2A) and *Thor* (4E-BP1 ortholog): (2) receptor kinase signaling elements, like *branchless, eiger* (TNFₗ ortholog), *Star* (EGFR ligand transporter), and *pointed* (ETS transcription factor ortholog): (3) regulators of chromatin and histone methylation, like *toutatis* (chromodomain protein, *kismet,* MEP-1, *windei* (H3K9 methyl transferase) and *Kdm2* (lysine specific demethylase). Thus, new LexA enhancer trap lines presented here significantly expand the arsenal of available LexA expression tools (Kockel et al., 2016, Pfeiffer et al., 2013).

Prior studies have demonstrated that P-element insertion in flies is non-random (O’Hare and Rubin, 1983; Berg and Spradling, 1991, Bellen et al, 2011), with a strong bias for transposition to the 5’ end of genes (Spradling et al., 1995). Here and in prior work, we have found a similar preference with *StanEx*^1^ P-element transposition; 89% of unique insertions were located in the promoter or 5’ UTR regions of genes. However, we also detected an unexpectedly high rate (9/192; 4.7%) of transposase-dependent *StanEx*^1^ P-element transpositions to a defined site on chromosome III within the *NK7.1/Heatr2* locus, at 3R:14,356,561. Subsequent analysis provided strong evidence for P element replacement of an undiscerned *KP* element at 3R:14,356,561 by the *StanEx*^1^ donor P-element as the basis of this finding **(Figure 3).** P-element ‘replacement’ as a mechanism explaining biased insertion frequency in our screens is supported by our findings of (1) concomitant *KP* deletion upon StanEx P-element insertion into 3R:14,356,561, (2) precise substitution into the 5’ and 3’ breakpoints defined by the prior *KP* element, (3) absence of 8bp direct repeats generated *de novo* by transposon-mediated integration of the *StanEx*^1^ element, (4) high rate of P-element insertion into the site occupied by the prior *KP* element, and (5) the not-yet-explained tendency of the donor P-element to maintain the directionality of insertion of the outgoing P-element (Williams, 1988, Heslip and Hodgetts, 1994, Gonzy-Treboul et al., 1995, de Navas et al., 2014). *KP* elements in the genome of wild strains of *Drosophila melanogaster* have been reported frequently (Itoh et al., 2007), but the origins of this particular *KP* element remains enigmatic. In our search to identify the source of the *KP* element at 88B4-6, we screened eleven fly strains obtained from the Bloomington Stock Center and identified one strain of eleven, *w[l]* (stock 2390), that also harbored this *KP* element. Stock 2390 was added to the Bloomington Center in 1989 and might therefore not represent a true copy of the original *w[l]* strain (Morgan, 1910; Johnson, 1913).

Since the inception of P-element mutagenesis screens, hot spots for P-element insertions have been noted (Bellen et al., 2011). Prior enhancer trap screens, which initially used ⍰132-3 transposase-dependent strategy, later adopted an alternative ⍰2-3 transposase-independent approach (e.g., *PiggyBac;* Thibault et al., 2004, Gohl et al., 2011), due to unacceptably high insertional hot spot over-representation (Bellen et al., 2011) and mobilization of undetected *KP* elements (Dr. T. Clandinin, personal communication). In these prior screens, it remains unknown if an occult *KP* element also distorted insertion site selection and frequencies. Our findings suggest that *KP* elements can contribute to phenotypes like insertional over-representation following transposase-mediated enhancer screens. However, in practice, the *StanEx* insertion bias to 3R:14,356,561 did not impact the productivity or strategy of our screen.

The mutated transposase ORF in *KP* elements is thought to encode an inhibitor that can suppress *P*-element transposition (Rio, 1990; Rio, 1991; Simmons 2016). This suppression has been suggested to be *KP*-dose dependent (Sameny and Locke, 2011), or caused by *KP* elements with distinct chromosomal location and high inhibitory activity (Fukui et al., 2008). The *KP-* element observed in this study is located at 88B4-6 and has not been previously described (Fukui et al., 2008). While the rate of *StanEx*^1^ transposition in the absence of the *KP* at 88B4-6 is unknown, the data here and our prior work (Kockel et al 2016) show that *StanEx*^1^ element transposition was not overly inhibited by this *KP* element.

The resources and outcomes described here significantly extend and develop the interscholastic partnership in experiment-based science pedagogy described in our prior study (Kockel et al 2016), which involved researchers at Stanford University and a single biology class at an independent secondary school. A curriculum based on fruit fly genetics combined with developmental and molecular biology provided an ideal framework for offering authentic research experiences for new scientists, including practical and curricular features detailed in Table 1 and prior work (Kockel et al 2016; Redfield, 2012). These course features offered both students and teachers a tangible prospect of generating one or more novel fly strains, thereby promoting a sense of discovery and ownership (Hatfull et al., 2006), and connection to a broader community of science, each key research and educational goals. For example, within the first 9 months of submission of StanEx lines described in our prior work to the *Drosophila* Bloomington Stock Center there were 153 strain requests from 63 labs in 15 countries. Use of *StanEx*^1^ lines, in publications and through direct requests (e.g., Babski 2018; Babski et al 2018; Cohen et al 2018; Drs. L. O’Brien, A. Baena-Lopez, personal communication) are additional indicators of practical outcomes from our work.

**Table 1:**
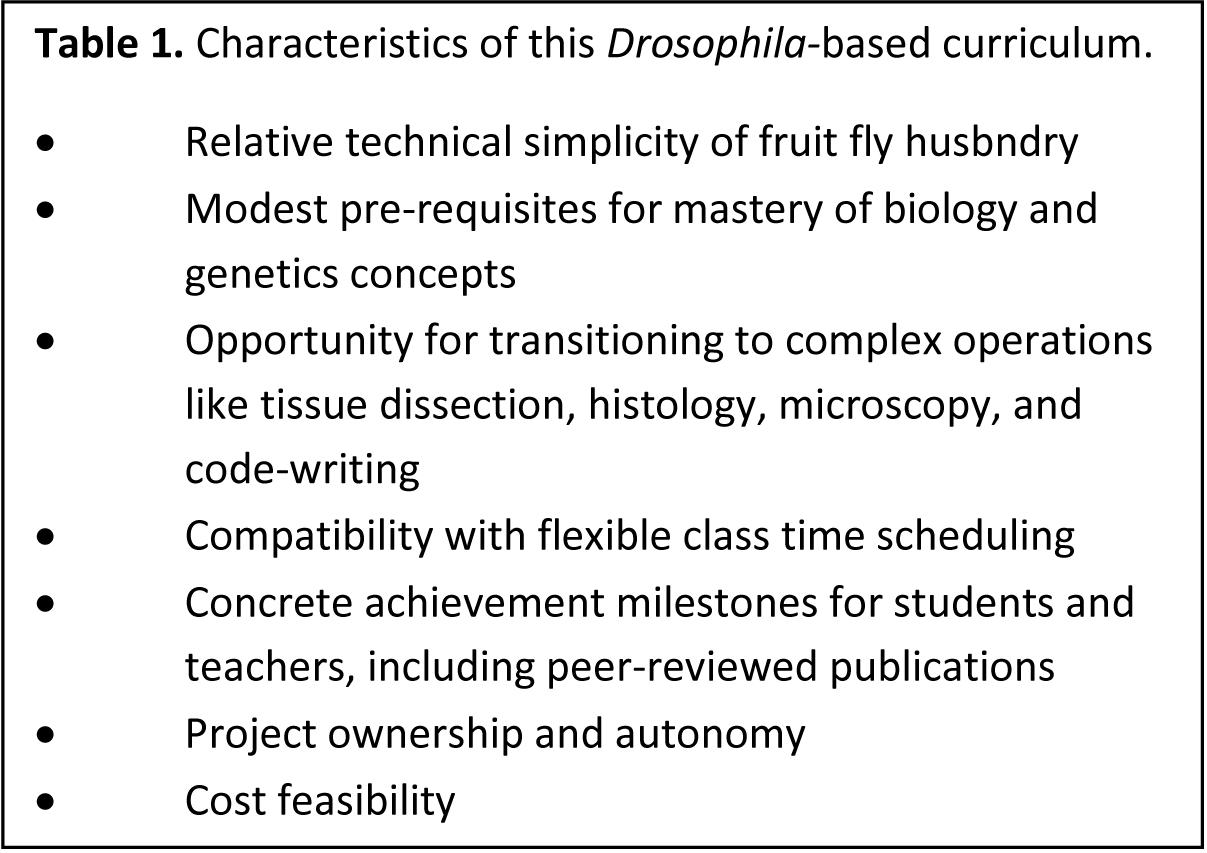
Characteristics of this *Drosophila*-based curriculum. Goals of course, expectations towards students, and the relative difficulty to achieve them as judged by the authors are listed. See text and (Hatfull et al., 2006) for details.

Here, we show that that our interscholastic partnerships and classroom-based research have expanded to include high schools in four states in the U.S.A. These schools encompass a spectrum of public, charter, independent and ‘high needs’ schools, with day or boarding students. This expanded framework has fostered additional curricular attributes, including (1) development of student leaders, who teach peers and novice adult teachers, and develop curricular innovations, (2) interscholastic collaboration and data sharing through regular video conferencing organized by university researchers, (3) additional professional development for adult teachers who train or mentor novice incoming adult teachers, and create new course content tailored to their student body, (4) infrastructure development to accommodate course expansions (like larger and/or additional courses: **Figure 4),** and **(5)** improved placement of student course graduates in competitive university- and industry-based summer research internships. In summary, this experience demonstrates how longitudinal studies involving multi-generational genetics, molecular and developmental biology, and bioinformatics developed by university researchers can build a thriving, interconnected network of secondary school teachers, students and classes that impacts science, personal growth, and professional development.

## Acknowledgments

We thank past and current members of the Kim lab for helpful discussions and welcoming summer term students. We are grateful to Philip Weissman (Micro-Optics Precision Instruments, NY) and Ken Fry (Genesee Scientific, CA) for generous support of equipment procurement for this project, and Dr. Johannes Albrecht Birgmeier for the polynomial probability calculation. We thank Drs. Tom Clandinin, Don C. Rio and Jeff Sekelsky for their insightful comments. We thank Stephen Murray, Greg Hansen, liana Saxe (The Lawrenceville School), Alison Hobbie, John Blackwood, Pamela Parris, Ethan Shapiro, Thomas Hassan, Lisa McFarland, and William Rawson (Phillips Exeter Academy), Andrea Beatty, Alison Offerman-Celentano and Jeanne Suttie (Commack High School), Carrie Spitz and Paige Moran (Pritzker College Prep) for their advice, support and encouragement. Work at Phillips Exeter Academy was supported by the John and Eileen Hessel Fund for Innovation in Science Education. We thank Glenn and Debbie Hutchins, and the Hutchins Family Foundation, for supporting opportunities for students to engage in innovative science research at the Lawrenceville School.

## Supplemental Figure Legends

**Suppl. Table 1:** Molecular data associated with StanEx enhancer trap lines **(Figure 1).** The data is also available in a searchable format at the StanEx online database http://stanex.stanford.edu/about/. The molecular insertion coordinate is defined as the first nucleotide 3’ of the genomic scaffold *independent* of the direction of the *P*-element insertion.

**Suppl. Table 2:** Molecular sequence of the 14 bp insertion site of 128 StanEx enhancer trap lines **(Figure 3A).** The sequences are oriented according to the strand of insertion. P-elements inserted in 5’->3’ direction are marked + and are co-directional with the genomic scaffold of the reference sequence. P-elements inserted in 3’->5’ direction are marked - and the reverse complement of the genomic scaffold sequence is used.

**Suppl. Table 3:** Chi-square *p*-values of position frequency matrices of StanEx enhancer traps **(Figure 3, Suppl. Table 3)** versus position frequency matrices of EPgy2, GT, SUPor-P, XP and GawB(+) enhancer traps as well as their agglomerate. Associated worksheets present primary sequence information and position frequency matrices of individual types of enhancer trap elements and is based on (Linheiro and Bergman, 2008).

**Suppl. Table 4:** Molecular data of the insertion site in the *NK7.1/Heatr2* locus at 3R:14,356,562 88B4-6 on chromosome III. Worksheets show (1) name and sequences of primer used to amplify genomic DNA from StanEx^1^ background, (2) genomic sequence according to the reference sequence in Flybase (Hoskins et al., 2015), (3) genomic sequence of StanEx^1^ background, (4) KP element sequences present in Genbank, (5) KP element sequence of the locus *NK7.1/Heatr2* locus at 3R:14,356,562 88B4-6, including genomic sequence 5’ (black) and 3’ (blue) to KP element, 8bp direct repeat (red) Exon (light/dark brown, grey)/lntron (black) structure of KP repressor encoded by KP element, KP repressor cDNA, amino-acid sequence of KP repressor protein translated from cDNA *in silico,* GenBank accession number MK510925 (6) Sequence data of SE enhancer trap lines inserted in 88B4-6, 3R:14,356,562 **(Figure 3C).**

## Supplementary Results

The canonical insertion of a P-element into a target site is marked by the emergence of a direct duplication of an 8 bp sequence flanking the 5’ and 3’ end of the P-element point of incision (Position number marked in orange, **Figure 3A and 3B**, O’Hare and Rubin, 1983). Therefore, the substitution of one P-element (the KP element) with another (the StanEx P-element) could leave two sets of 8bp target site duplications when the excision - insertion occur in series, and only one set when the events occur concomitantly. To distinguish between these two models, we analyzed the flanking genomic sequence of the nine StanEx insertions at the 3R:14,356,561 site for direct repeats and compared these sequences to the direct repeat flanking the KP insertion. The only 8 bp direct repeat detected in all cases, KP as well as all StanEx insertions at 3R:14,356,561, was GCC CAA CC. No other additional sequence duplication was noted at this site in all StanEx insertions. The data therefore supports a model of concomitant excision of the KP combined with the insertion of the StanEx element.

The exact replacement of one P-element by another without the generation of an additional 8 bp target site duplication has been observed before (Williams, 1988, Heslip and Hodgetts, 1994, Gonzy-Treboul et al., 1995, de Navas et al., 2014). These events have been reported to occur frequently (4-30%) and were aptly named P-element replacement. While a preference towards conserved P-element directionality has been reported during P-element replacement (Gonzy-Treboul, 1995), inversions of direction of the replacing P-elements have been observed (Heslip and Hodgetts, 1994, De Navas, 2006). In this study, we report two out of the nine cases of P-element replacement with inversed direction, SE457 and RJ-3 **(Figure 3C)**.

P-element replacement predicts the conservation of genomic sequences 5’ and 3’ of the insertion site. In some cases of P-element replacement, however, deletions, rearrangements and duplications of genomic sequences during P-element replacements have been reported (Gonzy-Treboul, 1995). To assess the status of adjacent genomic DNA, we sequenced the genomic DNA 5’ and 3’ of the breakpoint of the nine P-element replacements at 3R:14,356,561 (Methods). Out of the nine insertions, seven were conservative and did not show changes of 5’ or 3’ genomic sequence. Two lines, SE304 and SE405, did display alterations of genomic DNA 5’ to the insertion site. SE305 exhibits a nine base pair deletion of the 5’ 8bp direct repeat, and the first nucleotide of the 5’ end of the StanEx P-element. SE405 presents a 31 bp deletion of the 5’ P-element end and a further deletion/insertion of genomic sequences 5’ to the insertion site **(Suppl. Table 4)**.

